# Stem canker caused by *Phomopsis spp*. induces changes in polyamine levels and chlorophyll fluorescence parameters in pecan leaves

**DOI:** 10.1101/2021.03.04.433926

**Authors:** Guillermo Martin Mantz, Franco Ruben Rossi, Pablo Esteban Viretto, María Cristina Noelting, Santiago Javier Maiale

**Affiliations:** Instituto Tecnológico de Chascomús (INTECH), Consejo Nacional de Investigaciones Científicas y Técnicas (CONICET)-Universidad Nacional de San Martín (UNSAM), Int. Marino Km 8, Chascomús, Provincia de Buenos Aires, Argentina; Estación Experimental Agropecuaria Valle Inferior del Río Negro (EEA)-Instituto Nacional de Tecnología Agropecuaria (INTA), Valle inferior Río Negro, RN 3 Km 971, Pcia. RN, Argentina; Instituto Fitotécnico de Santa Catalina (IFSC), Universidad Nacional de La Plata (UNLP), Garibaldi 3400, Lavallol, Provincia de Buenos Aires, Argentina

**Keywords:** pecan, *Phomopsis*, polyamines, chlorophyll fluorescence, proline

## Abstract

Pecan plants are attacked by the fungus *Phomopsis spp.* that causes stem canker, a serious and emerging disease in commercial orchards. Stem canker, which has been reported in several countries, negatively affects tree canopy health, eventually leading to production losses. The purpose of this study was to inquire into the physiology of pecan plants under stem canker attack by *Phomopsis spp*. To this end, pecan plants were inoculated with an isolate of *Phomopsis spp.* and several parameters, such as polyamines, proline, sugars, starch, chlorophyll fluorescence and canopy temperature were analysed. Under artificial inoculation, a high disease incidence was observed with symptoms similar to those in plants showing stem canker under field conditions. Furthermore, the infected stem showed dead tissue with brown necrotic discolouration in the xylem tissue. The free polyamines putrescine, spermidine, and spermine were detected and their levels decreased as leaves aged in the infected plants with respect to the controls. Chlorophyll fluorescence parameters, such as Sm, ψEO, and QbRC decreased under plant infection and therefore the K-band increased. Canopy temperature and proline content increased in the infected plants with respect to the controls while sugar content decreased. These data suggest that stem canker caused by *Phomopsis spp.* induces physiological changes that are similar to those observed in plants under drought stress. To our knowledge, this is the first study that documents the physiological and biochemical effects derived from pecan-*Phomopsis* interaction.

## 1 INTRODUCTION

The pecan tree (*Carya illinoinensis* (Wangenheim) K. Koch) produces seeds with high nutritional value and medicinal properties (Atanasov *et al*., 2017). It is native to North America (USA and Mexico) and is cultivated not only in the area of origin but also in many other countries in the temperate zone. Pecan plants are continuously exposed to different environmental stresses like pathogen attacks which cause a significant economic impact in the agricultural industry and lead to yield losses. Pecan canker was originally detected in Pima County (Arizona, USA) and was reported to be caused by *Cytospora spp.* (Hine *et al*., 1969), affecting scaffolds and other branches in two- and three-year-old plants. In this first approximation, Hine *et al*. observed that superficial lesions, such as those caused by low winter temperatures or sunburn in summer, are necessary to allow the fungal pathogen to enter host tissues and concluded that disease severity is genotype-dependent. Two types of Shoot Dieback Maladies (SDM) of pecan were further found to adversely affect tree canopy health in early spring and early summer in the USA pecan belt region. *Phomopsis spp.* and *Botryosphaeria spp.* were found to be the causal agents of SDM in 14 cultivars evaluated, representing 89% and 40% of the isolates, respectively (Reilly *et al*., 2010). Decline and mortality of hickory species belonging to the *Carya* genus (e.g., Shagbark and Bitternut) have been linked to *Ceratocystis smalleyii*, *Fusarium solani*, and *Phomopsis spp.* (Juzwik *et al*., 2008). The symptoms reported by Juzwik *et al*. included sunken small stem canker associated with *Ceratocystis* and *Fusarium* species and galls on the stem and branch associated with *Phomopsis spp*. Of the isolates documented in Juzwik *et al*.’s study, four were morphologically identical to those described as *Phomopsis spp.* and two of them matched in the GenBank database with *P. amygdale* and *Diaporthe helianthi*.

In Argentina, several diseases are known to affect the growth and development of pecan orchards, such as scab caused by *Venturia effusa* (ex-*Cladosporium caryigenum*) (Mantz *et al*., 2008), anthracnose on the pecan shuck and leaves caused by *Colletotrichum gloeosporioides* (Mantz *et al*., 2010) and nut pink mould caused by *Trichothecium roseum* (Mantz *et al*., 2009). All the latter have been recorded in the Pampas plains from central Argentina under temperate and moist weather conditions. A new disease whose causal agent was isolated from Pawnee cultivar has been recently recorded in Argentinean pecan orchards. It affects both the stems and branches and is typically characterised by canker symptoms. It was reported to belong to the genus *Phomopsis spp.* (Noelting *et al*., 2016).

*Phomopsis* is the asexual state of *Diaporthe* contains more than 900 species with broad hosts range and worldwide distribution, and the taxonomy of this fungus is constantly evolving (Udayanga *et al*., 2011, Gomes *et al*., 2013). Although *Diaporthe* is preferred to name this group of fungi, as in this work the anamorph was used, henceforth *Phomopsis* will be used to refer to this pathogen.

The symptoms include sunken and elongated lesions, particularly 0.5 to 1 cm long cankers mainly in the branches at the level of the neck or in the area of rootstock-scion union. These lesions are characterised by the presence of dark structures corresponding to fungus fruiting bodies (pycnidia). Cankered plants are characterised by bark sloughing and twig or branch death. These symptoms have been observed with high incidence and severity in numerous cultivars, such as Colby, Starking hardy giant, Desirable, Stuart, Lucas, Hirschi, and Pawnee, in the southern Argentinean Pampean region (Mantz, personal observation).

Under pathogen attack, plants use a wide variety of physical and chemical barriers derived from infection (Grant and Lamb, 2006). Polyamines (Pas) are one of these chemical barriers related to plant defence. They are natural aliphatic polycations essential for most organisms. At physiological pH, Pas are positively charged and therefore interact with anionic molecules, such as proteins, phospholipids, and nucleic acids. Diamine putrescine (Put), triamine spermidine (Spd), and tetramine spermine (Spm), which are the most abundant polyamines in plants (Bagni and Tassoni, 2001), are molecules involved in key processes, such as growth, development, morphogenesis, embryogenesis, senescence, and response to abiotic and biotic stress like plant defence mechanisms (Kusano *et al*., 2008). During plant-microbe interaction, Pas undergo a remarkable change (Jiménez Bremont *et al*., 2014, Romero *et al*., 2018). In line with this, previous research has shown that Pas content and metabolism are augmented in plants under pathogen attack independently of whether the pathogen infection strategy is via biotrophic or necrotrophic interactions (Walters, 2003; Jiménez Bremont *et al.* 2014; Romero *et al*. 2018). It has also been demonstrated that phytopathogens perturb the activity and functionality of photosystem and Pas levels (Vilas *et al*., 2018). Plants respond to biotic stress by adjusting their machinery to maintain photosynthetic activity. Once stress has overcome the acclimation capacity, permanent photoinhibition and inhibition of the electron transport chain occur (Perez-Bueno *et al*., 2019). Evidence from previous research (Hamdani *et al*., 2011) suggests a connection between Pas metabolism and photosynthetic activity. Exogenously added Spm may penetrate into the thylakoid membranes and interact with proteins and extrinsic polypeptides of the oxygen-evolving complex (OEC). Consequently, PSII activity could be preserved and protected from photoinhibition under high light stress by Spm (Hamdani *et al*., 2011). In line with this, other authors found that Put is a stimulator of ATP biosynthesis while Spd and Spm are efficient stimulators of non-photochemical quenching (Ioannidis and Kotzabasis, 2007).

Taking all the above into account, the purpose of the present study was to inquire into the physiology of pecan under stem canker attack by *Phomopsis spp*. To this end, different physiological parameters, such as Pas levels, canopy temperature, and chlorophyll fluorescence were analysed. Results from these analyses are −to our knowledge− the first in reporting the physiological changes that occur in pecan leaves as a result of infection with *Phomopsis spp*.

## 2 MATERIALS AND METHODS

### 2.1 Biological material and growth conditions

Experiments were carried out under ambient conditions at the campus of the *Instituto Tecnológico de Chascomús* (INTECH), a research center depending on the National Research Council (CONICET) and the Universidad Nacional de San Martín (UNSAM) in Argentina. Meteorological data were taken from the Automatic Weather Station located at the Experimental Station “Manantiales”, a dependency of the *Instituto Nacional de Tecnología Agropecuaria* (INTA) and the *Ministerio de Asuntos Agrarios* in Buenos Aires province, Argentina (Supplemental Figure 1). The pecan plants, cv. Pawnee, analysed herein were obtained from a local nursery, and the cultivar is the same as that where the pathogen used in this study was isolated (Noelting *et al*., 2016). The plants used as rootstock were produced from the pecan nut obtained from local plant seedlings and they were grown in a 10 L pot for two years. Rootstocks were grafted in autumn with buds from Pawnee, the scion was cultivated for one growing season, and the following year they were used for the experiments herein described. Control and inoculated plants were placed in a plastic pool and sub-irrigated with rainwater twice a week.

*Phomopsis spp.* isolate number 1219, which corresponds to the isolate described by Noelting *et al*. (2016), was obtained from the *Instituto de Botánica “Carlos Spegazzini”* belonging to the *Universidad Nacional de La Plata*, in Argentina. The fungus was routinely maintained on PDA agar at 22 °C with a 12 h photoperiod to induce conidia generation and was sub-cultured biweekly.

### 2.2 Plant inoculations

Inoculation was performed on 18 plants following Noelting *et al*. (2016). Briefly, in November 2018, plugs of 0.5 cm in diameter of *Phomopsis spp.* growing actively on PDA were used to inoculate pecan plants. Shoots growing during the year of the experiment were used for inoculations, a 1 cm cut was made above the petiole insertion with a scalpel and a mycelial plug was placed on the lesion. A piece of cotton wool moistened in sterile water was placed on the mycelia plug to prevent mycelial dehydration. Polyethylene film was used to hold the plug closely attached to the petiole. Control treatments were carried out on other 18 plants following the same procedure and on the same day as for the infected plants but using a PDA disk without mycelium.

### 2.3 Free polyamine quantification

Samples for Pas quantification were harvested on days 9 and 23 after inoculation (DAI). At each time point, the fully developed leaflets of leaves located above the inoculation point were detached and stored at −18 °C until use. Polyamines were analysed by derivatisation with dansyl chloride, separated in HPLC reverse phase, and detected by fluorescence according to Maiale *et al*. (2004) with modifications. Briefly, samples were powdered using a mortar and pestle precooled with N_2_ liquid. Three hundred milligrams of plant powder were placed in a 1.5 ml microtube, and 1 mL of 5% HClO_4_ was added, vortexed, and placed on ice for 30 min. Samples were subsequently centrifuged at 10000xg for 10 min, 60 μL of the sample were transferred to a microtube containing 60 μL of saturated Na_2_CO_3_, 6 μL of 0,1 mM 1,7 heptanodiamine (Htd) as internal standard, and 75 μL of 10 mg/mL dansyl-chloride in acetone. Samples were incubated at 70 °C for two hours in the dark, the reaction was stopped by adding 20 μL of L-proline (100 mg/mL) and incubated for 15 min under the same conditions. Dansylated Pas were extracted with 200 μL of toluene, vacuum dried, and kept at −18 °C until quantification.

A HPLC binary pump (Waters 1525) attached to a C18 reverse-phase column 25x 4 mm Luna (2) (Phenomenex) and a diode array fluorescence detector (Waters 2475) were used in gradient protocols with acetonitrile (Act) and H_2_O. Gradient protocols were used with a total time of 15 min following these steps: 0 to 4.5 min Act 70% - H_2_O 30%, 4.5 to 5.5 min shifted to 100% Act, 5.5 to 9 min hold to 100% Act, 9 to 10 min shifted to Act 70% - H_2_O 30% and 10 at 15 min maintain Act 70% - H_2_O 30%. Dried samples were dissolved in 40 μL acetonitrile and injected in the HPLC through a 5 μL loop capacity. The fluorescence signal was detected at 415 nm excitation and 510 nm emission and integrated and processed using Breeze^®^ software (Waters). A calibration curve with synthetic Pas was performed and Pas levels were calculated.

### 2.4 Chlorophyll fluorescence fast-transient analysis

Chlorophyll fluorescence was performed with a portable fluorometer (HandyPEA, Hansatech Instruments, UK) on 0, 9, 16, and 23 DAI following Corigliano *et al*. (2019), with modifications. Central leaflets of intact leaves above the inoculation point were pre-darkened for 20 min before analysis using a leaf clip provided by the manufacturer and they were subsequently exposed during 1 s to 3000 μmol photons m^−2^s^−1^ (650 nm peak wavelength) with a dark interval of 500 ms and exposed during 1s to 3000 μmol photons m^−2^s^−1^ again and chlorophyll-a fluorescence was recorded. The fluorescence data were processed by PEA plus software (Hansatech Instruments, U.K.) to obtain OJIP parameters. A summary of the OJIP parameters used in this study is shown in Table 1.

**Table 1:**
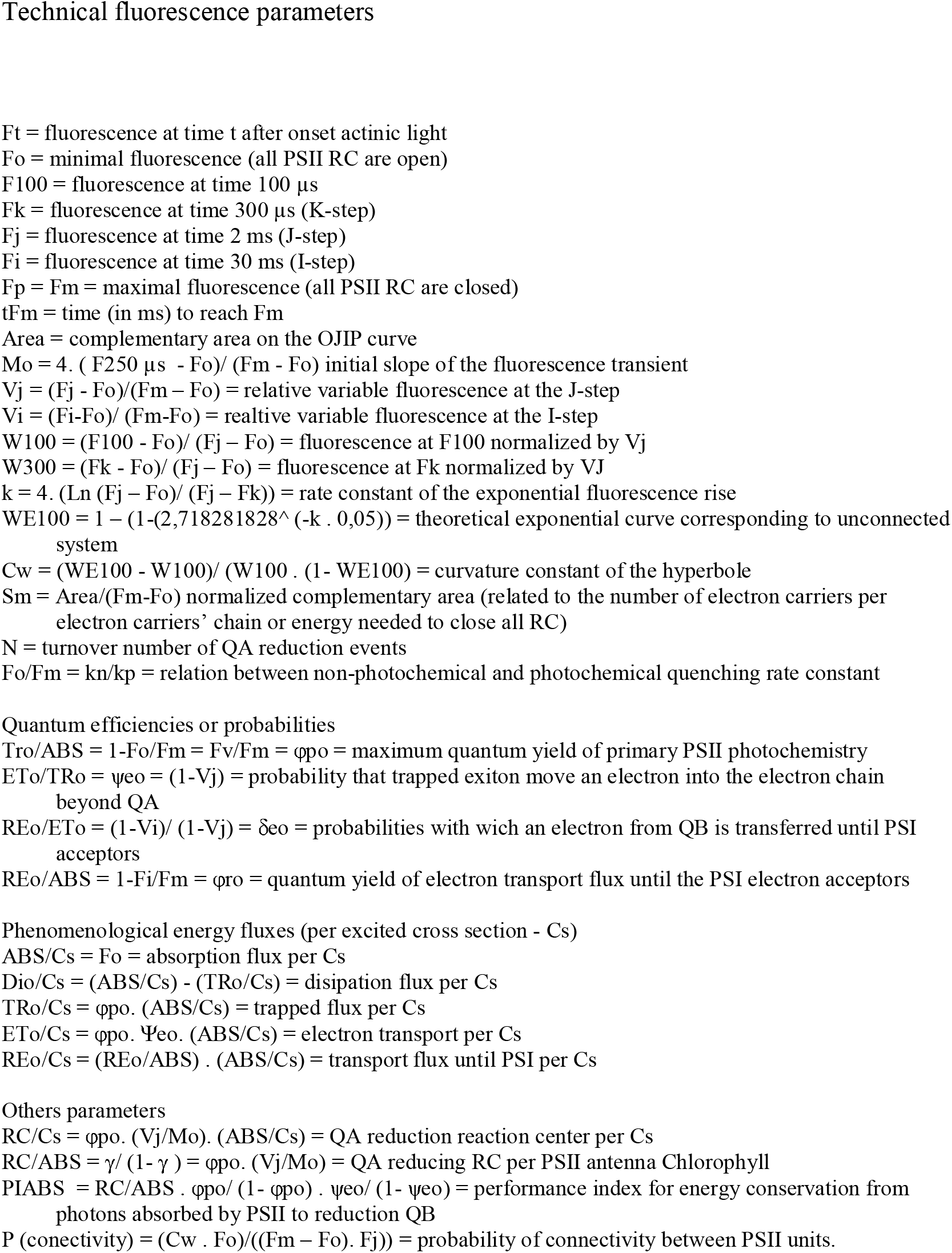
Chlorophyll fluorescence parameters analysed by OJIP test.

### 2.5 Canopy temperature measurements

Canopy temperature was measured according to Rachoski *et al.* (2015) on 6, 10, 16, and 32 DAI using a thermal camera E-30 (FLIR, USA) with a resolution of 160 x 120 pixels and a thermal sensitivity of 0.1 °C. The camera was calibrated at 2 metres, 20 °C ambient temperature, 80% relative humidity, and an emissivity of 0.98. Thermographic pictures were analysed with the ThermaCam Research PRO (FLIR, USA) software.

### 2.6 Free sugar, starch, and proline determination

Free sugars and starch were determined using the anthrone assay according to Corigliano *et al*. (2019) with some modifications. Leaflets were homogenised in liquid nitrogen. Sugars were extracted by adding 1 mL of EtOH 80% v/v to 10 mg of powder and heated at 85 °C for 30 min. The samples were subsequently centrifuged at 10000 rpm for 10 min, and the supernatant was transferred to a clear tube. This procedure was repeated three times and the supernatant was pooled. The pellet was dried and 100 μL of HClO_4_ 35% were added and shaken in a vortex for 15 min and centrifuged for 10 min at 10000 rpm for starch solubilisation. Ethanolic or perchloric extract (6 μL) was mixed with 54 μL of distilled water, 240 μL of anthrone reagent (200 mg/mL in H_2_SO_4_ 72%), and incubated at room temperature for 5 min. The total sugar and starch content was calculated by reading the absorbance at 630 nm in a microplate reader (BioTek Synergy H1. VT, USA) using a calibration curve performed with glucose.

Proline content was determined following Babuin *et al*. (2016). Briefly, powdered leaflet samples (250 mg FW) were placed in a 2 ml microtube and boiled in 1 mL of distilled water for 30 min. Samples were centrifuged at 10000 rpm for 10 min, 180 μL of supernatant were transferred in a 2 mL microtube containing 180 μL of sodium citrate buffer (0.2 mol/L at pH 4.6) and 720 μL of 1% ninhydrin solution in acetic acid: water (60:40) and boiled at 96 °C for 1 h. Microtubes were subsequently cooled on ice, and proline was extracted with 720 μL of toluene by shaking in a vortex for 30 sec. The organic phase was read in a spectrophotometer at 520nm and a calibration curve with synthetic proline was constructed. Free sugar, starch, and proline content were determined on 9 and 23 DAI.

Dry weight was determined on 9 and 23 DAI by drying the leaflets in an oven at 70 °C and weighing up to constant weight. The biochemical data corresponding to Pas, proline, sugar, and starch were expressed on a dry weight basis.

## 3 RESULTS

### 3.1 Disease symptoms

The fungus strain used in this work was isolated from the pecan production zone of southern Argentina and its ITS sequence was previously deposited in the GenBank (accession N° 359871).

Using the Mycobank database, it was possible to observe that *Phomopsis spp.* showed high pairwise similarity values with two *Diaporthe spp.* species, reaching 99.42 in each case, and with *D. helianthi* and *D. infecunda*, reaching values of 99.226 and 99.156, respectively. A phylogenetic tree was constructed with a Mega 6.06 software (Supplemental Figure 2) but no clear information could be obtained, *D. helianthi*, *D. infecunda*, and *D. middletoni* being the most related species. Further thorough research studies are necessary to determine the correct taxonomy of *Phomopsis spp.*

Inoculation was carried out in spring under ambient conditions. Control plants showed no visible symptoms of disease throughout the experimental period whereas typical stem canker symptoms were observed in almost all the infected plants, except for three plants in which infection was not successful. The latter were therefore not considered in subsequent analyses. These symptoms consisted of tissue necrosis in the inoculation zone with formation of small cankers. Under the microscope, the transversal section of the shoot on 24 DAI above the infection point showed a dark brown colour in the xylem tissue, while control plants showed no symptoms (Figure 1A-B). Black pycnidia were observed on the dead tissue of the inoculated plant cankers (Figure 1D). The macroscopic symptoms and signs were morphologically identical to those already reported in pecan and different plant species infected with *Phomopsis spp*. (Uddin *et al*., 1997; Reilly *et al.*, 2010; Noelting *et al*., 2016).

**Figure 1:**
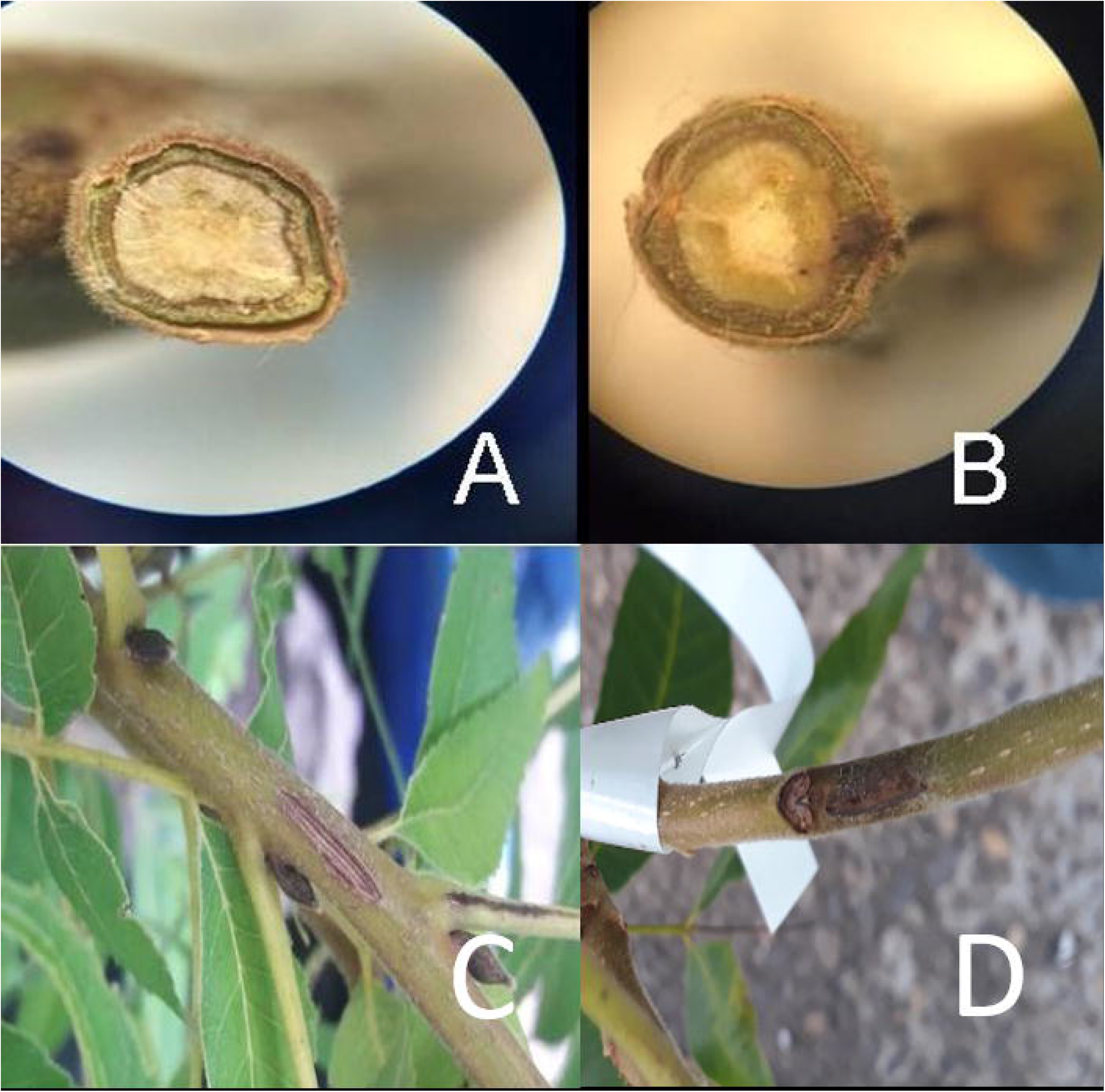
Transversal section of the stem above the point of infection on day 24 after inoculation and view of the area inoculated. A and C- mock-infected plant; B and D- plant inoculated with *Phomopsis spp*.

### 3.2 Changes in free polyamine levels

The changes that occurred in free Pas contents from pecan plants infected with *Phomopsis spp.* at different times were analysed. On 9 DAI it was observed that the most abundant Pas in the control pecan leaflets were Spd, followed by Spm and Put with 56%, 28%, and 16%, respectively (Figure 2 and Supplemental Figure 3) and that the levels of the three Pas detected in the infected plants were the same as those observed in the control ones. The levels of Spd and Put were observed to decrease over time in the control and infected plants, whereas Spm was found to remain constant. Put content on 9 DAI vs. 23 DAI showed a decrease of 68% and 81% in the control and infected plants, respectively. Spd content in the control plants on 9 DAI vs. 23 DAI underwent a decrease of 28% and 40% in the control and infected plants, respectively. Finally, Spm content in the control plants showed an increase of 19% while it maintained a constant value on 9 and 23 DAI. However, on 23 DAI Spm levels showed a significant decrease in the infected plants with respect to those in the control plants. Total Pas levels, therefore, showed a tendency to decrease over time, such a decrease being significant in the infected plants with respect to the controls (Figure 3).

**Figure 2:**
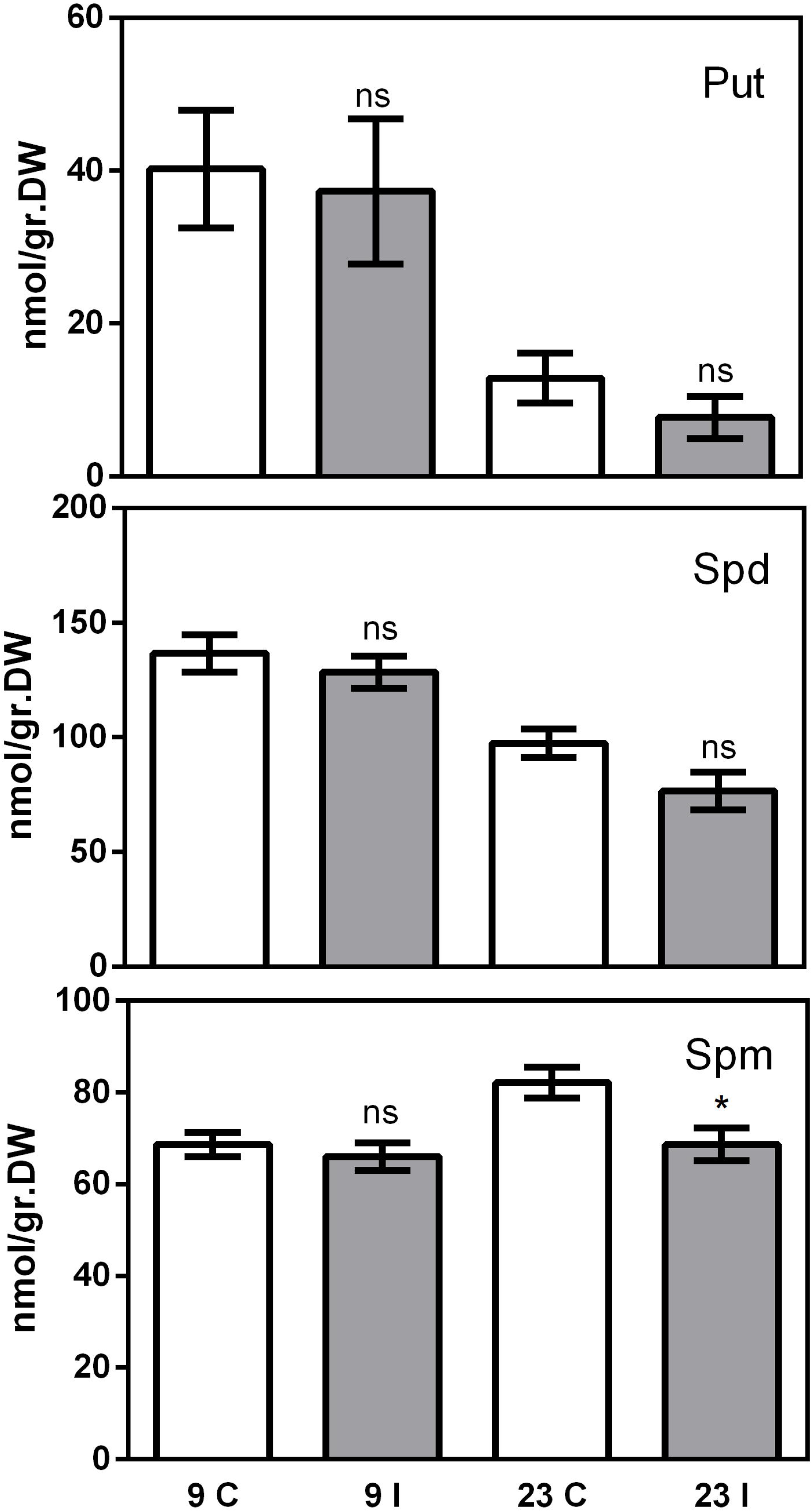
Polyamine content of putrescine (Put.), spermidine (Spd.), and spermine (Spm.) in control leaflets of pecan leaves (white bar) and in leaflets of pecan leaves infected with *Phomopsis spp.* (grey bar) on days 9 and 23 after inoculation. Data are mean with SEM and t-tests were performed for each sampling time comparing control plants and infected plants with a Graph Pad Prism^®^, ns-no significant; * p≤ 0.05; ** p≤ 0.01; *** p≤ 0.0001, n control: 16; n infected: 15.

**Figure 3:**
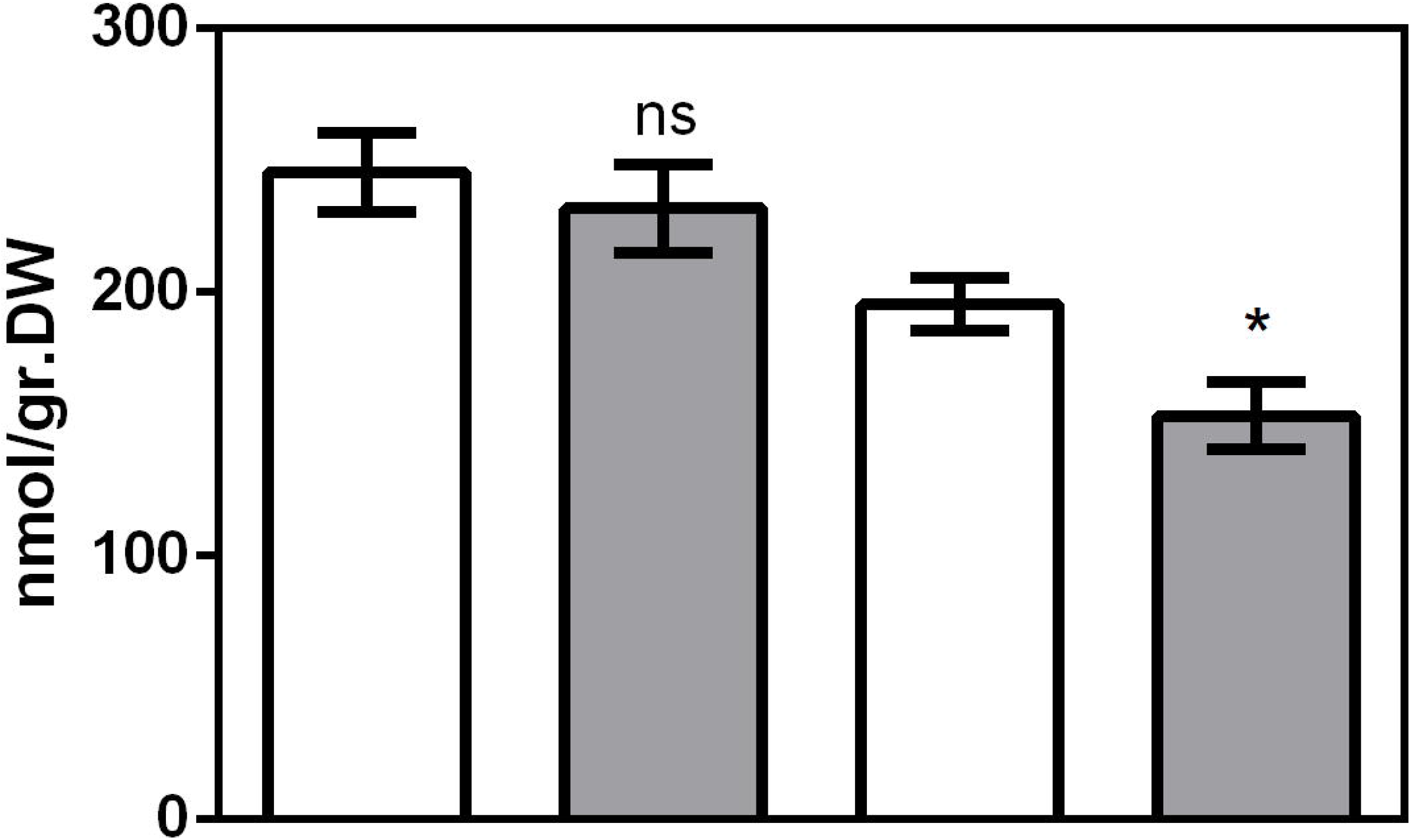
Total polyamine content in control (white bar) and infected (grey bar) pecan plants on days 9 and 23 after inoculation. Data are mean with SEM and t-tests were performed for each sampling time comparing control plants and infected plants with a Graph Pad Prism^®^, ns-no significant; * p≤ 0.05; ** p≤ 0.01; *** p≤ 0.0001, n control: 16; n infected: 15.

### 3.3 Measurement of chlorophyll a fluorescence transient parameters

Photosynthesis is an integral part of plant metabolism and is extremely sensitive to different stresses. In this respect, pathogenic infection often produces complex changes in the host□s photosynthetic apparatus which can be estimated by fluorescence measurements. In this work, fast transient chlorophyll a fluorescence was measured and an OJIP analysis was performed. Based on the data collected, normalised fluorescence curves were constructed. They showed a differential behaviour between the control and the infected plants. On 9 DAI the O-J and I-P steps of the infected plants showed differences with respect to those of the control plants, on 16 DAI the steps I-P and on 23 DAI the steps O-J showed a differential value between the control and infected plants (Supplemental Figure 4). At 0 time of inoculation, the control and infected plants did not show any significant differences in PIabs, Sm, Fv/Fm, and ψEO parameters. Nonetheless, these parameters were altered over the infection time course, and those particularly related to the health functionality of PSII were remarkably deteriorated (Figure 4E, 4F). Sm, ψEO, and PIabs parameters decreased along the experiment in the infected plants, yielding values of 88.9%, 91.3%, and 77.5% with respect to the controls on 23 DAI (Figure 4 B, 4C, and 4D). However, only ψEO and Sm showed a significant difference on 23 DAI.

**Figure 4:**
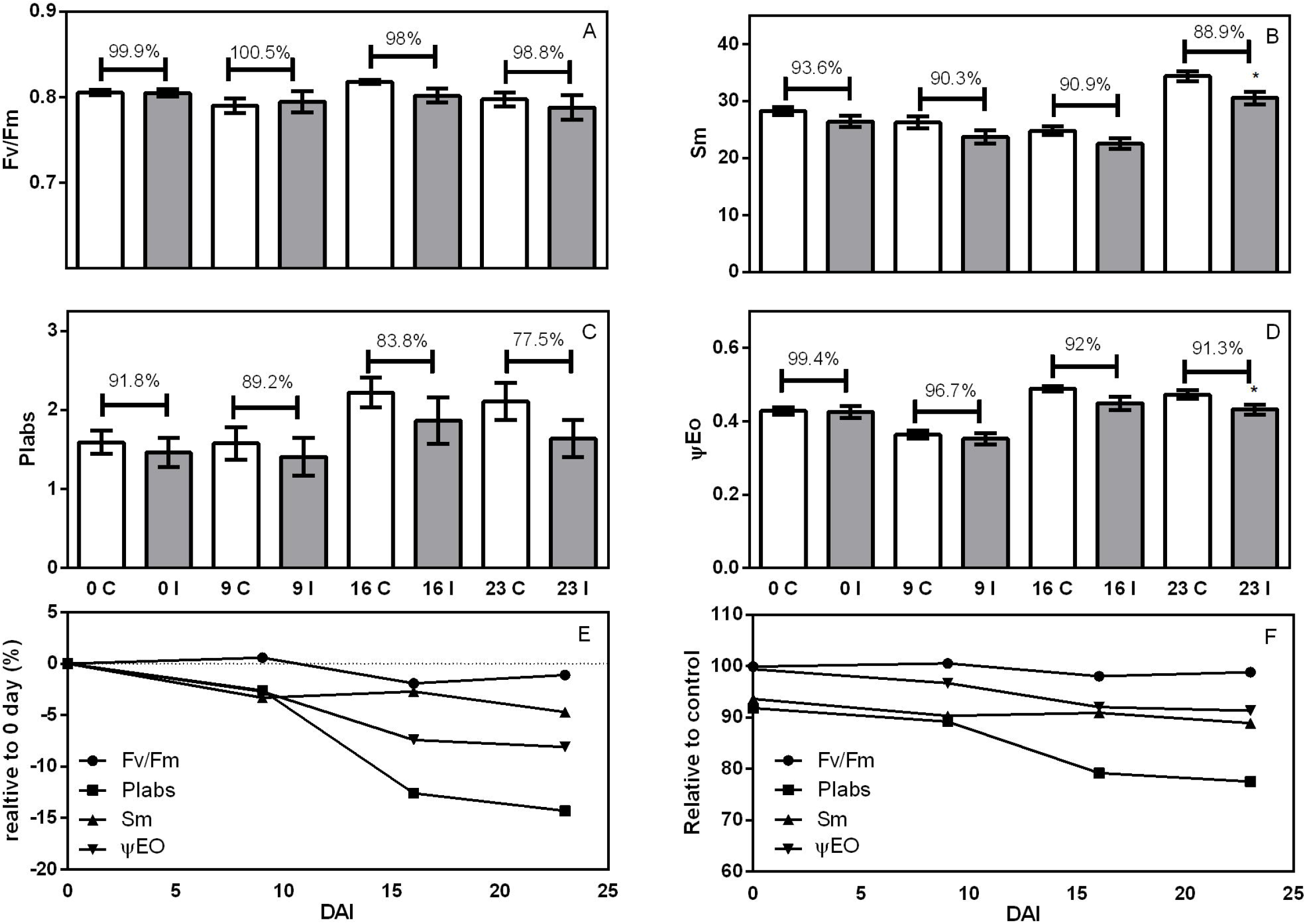
Chlorophyll a fluorescence parameters in leaflets of pecan control plants (white bar) and infected plants (grey bar) on days 1, 9, 16, and 23 after inoculation. A- Fv/Fm is the maximum quantum yield of primary PSII photochemistry; B- Sm is the electron transport chain per reaction centre; C- Piabs is a performance index for energy conservation from absorbed photons to reduction QB; D- ΨE_0_ is the probability that a trapped exciton moves an electron beyond QA; E- Fv/Fm, Sm, PIabs and ΨE0 values in percentage, normalised and compared with values in the control plants on day 0; F- Fv/Fm, Sm, PIabs and ΨE_0_ values in percentage compared with values in the control plants. Percentage in bars indicate the difference between the infected and control plants at each measurement time. Data are mean with SEM and t-tests were performed for each sampling time comparing control plants and infected plants with a Graph Pad Prism^®^, ns-no significant; * p≤ 0.05; ** p≤ 0.01; *** p≤ 0.0001, n control: 16; n infected: 15.

Other OJIP parameters, namely DIo/CSo, ABS/RC, DIo/RC, and TRo/RC, increased in the infected plants with respect to the control plants (Supplemental Figure 5). In addition, differential Woj showed an increase in the K-band between the infected and control plants along the experiments, and the RC fraction that reduced QB showed a significant decrease in the infected plants on 23 DAI with respect to the controls (Figure 5A).

**Figure 5:**
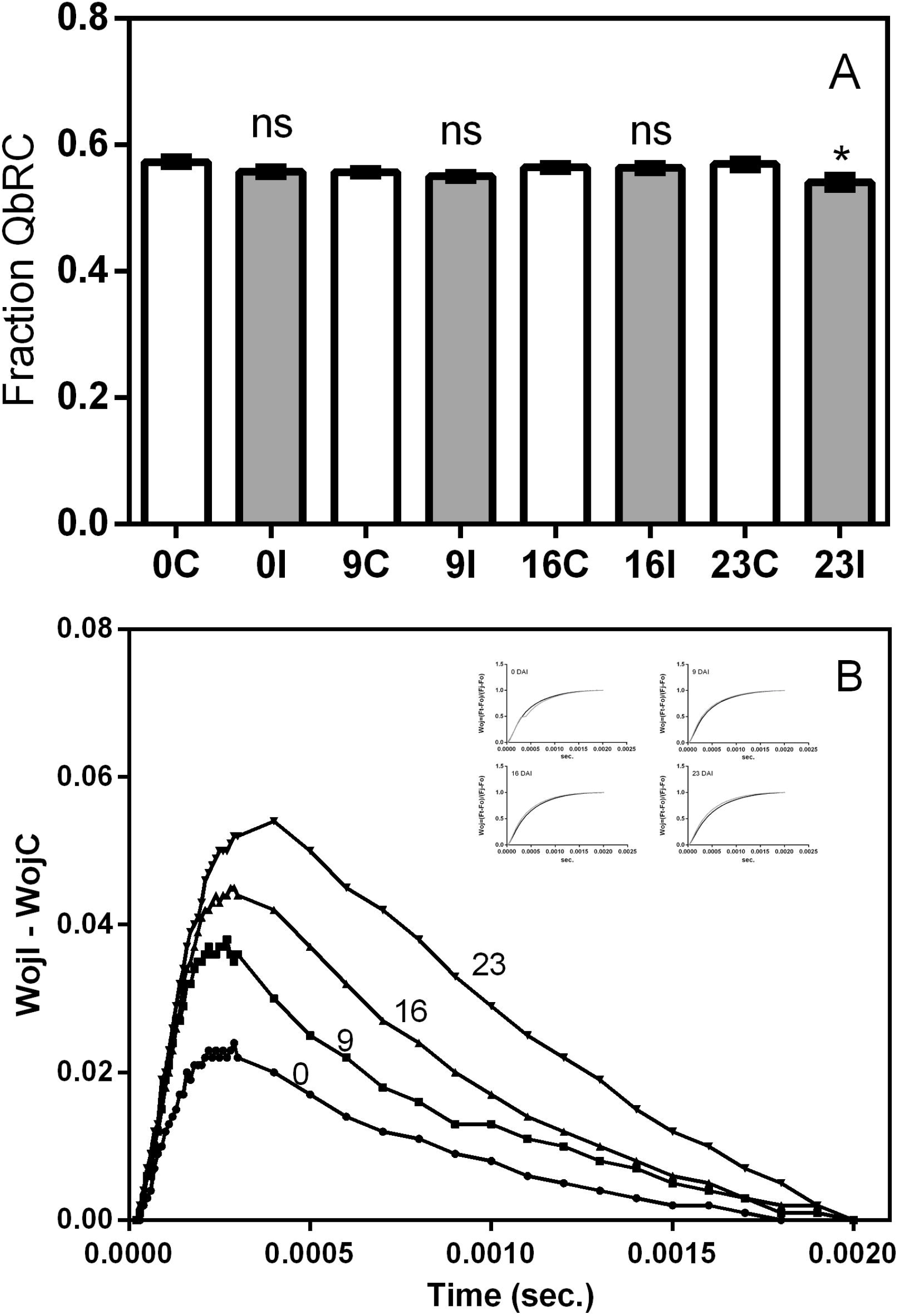
A- Fraction of the reaction centre that reduces QB in the controls (white bar) and infected plants (grey bar) on days 0, 9, 16, and 23 after inoculation. B- K-band (WojI-WojC) shown on days 0, 9, 16, and 23 after inoculation; insert Woj= (Ft-F_0_)/(Fj-F_0_) on the same day of measurement. Data are mean with SEM and t-tests were performed for each sampling time comparing control plants and infected plants with a Graph Pad Prism^®^, ns-no significant; * p≤ 0.05; ** p≤ 0.01; *** p≤ 0.0001, n control: 16; n infected: 15.

Furthermore, PIabs components, a multicomponent parameter related to energy conservation from photons absorbed by PSII to reduction QB (Table 1) showed a heterogeneous behaviour with RC/ABS tending to be less heterogeneous over time (Supplemental Figure 6).

### 3.4 Thermographic measurements of pecan plants

Leaf temperature is a consequence of the energy balance that includes the energy input of solar radiation and ambient heat and energy loss, such as, scattered light, heat loss, and transpired water. Pathogens may modify the values of these parameters when they infect plants (Bernard *et al*., 2013). For this reason, thermography becomes a useful tool to provide information on the transpiration of leaves attacked by a pathogen, especially in the early stages of pathogenesis when either symptom are not yet visible, or the infected area is small.

Plants inoculated with *Phomopsis spp.* showed an increase in canopy temperature which was measured with an infrared thermal camera (Figure 6). Although an increasing trend could −in fact− be observed in canopy temperature in the infected plants compared to the controls over the data collection time period in all the studied times (6-32 DAI), only on 16 DAI did the increase in temperature show significant differences.

**Figure 6:**
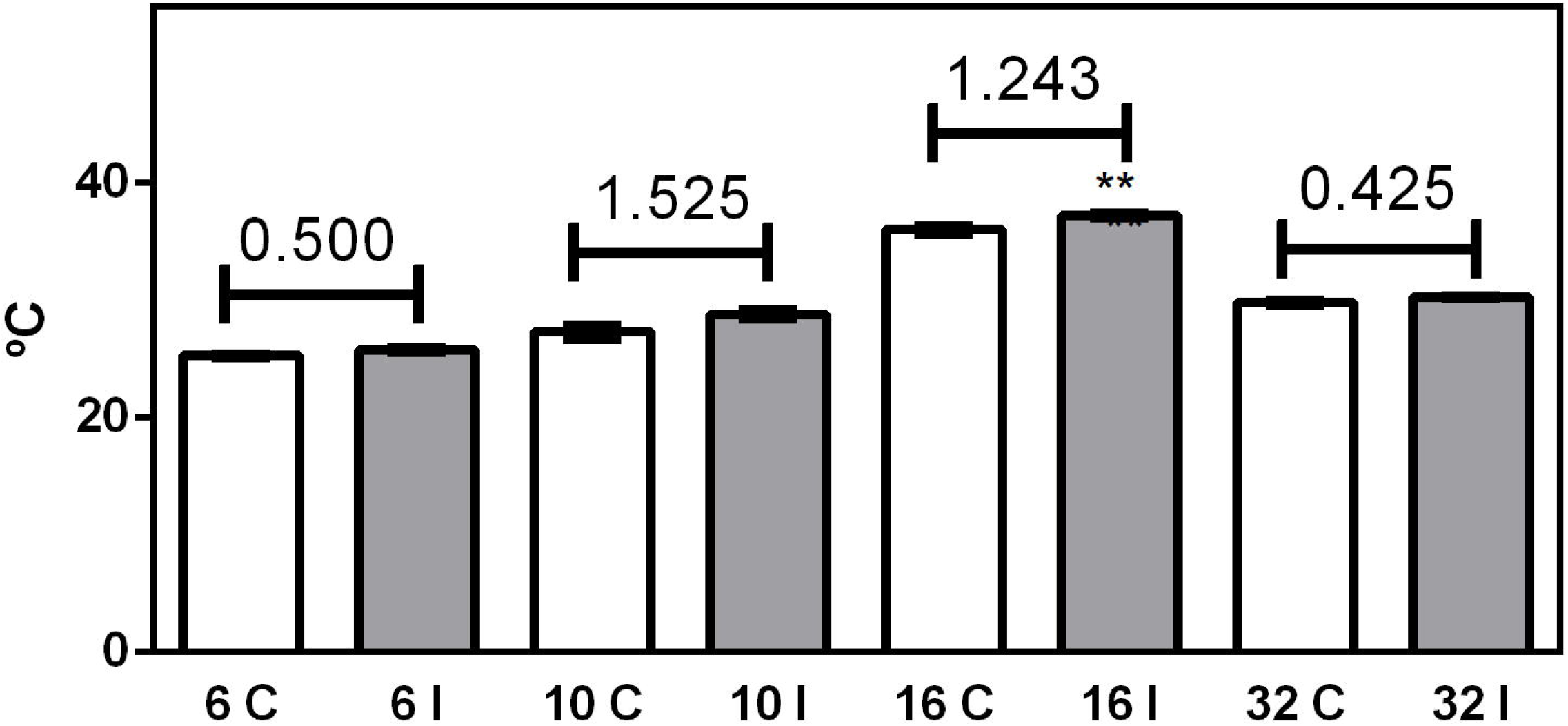
Canopy temperature in ° C measured by thermography on days 6, 10, 16, and 32 after inoculation in control plants (white bars) and infected plants (grey bars). Numbers above bars indicate differences in °C between infected and control plants. Data are mean with SEM and t-tests were performed for each sampling time comparing control plants and infected plants with a Graph Pad Prism^®^, ns-no significant; * p≤ 0.05; ** p≤ 0.01; *** p≤ 0.0001, n control: 16; n infected: 15.

### 3.5 Effects of *Phomopsis spp.* infection on free sugars, starch, and proline content

To study the levels of some organic solutes associated with different stressful conditions, free sugars, starch, and proline were measured in leaflets of pecan plants infected with *Phomopsis spp.* on 9 and 23 DAI. No significant differences were observed in free sugar or starch content at the two measurement times (Supplemental Figure 7). However, slightly higher levels of free sugars were observed in the control plants at both times analysed (17.42 and 14.19 vs. 16.95 and 13.25 mg of glucose eq. / 100mgDW in the control and infected plants, respectively) while starch values showed slightly higher values in the infected plants with respect to the controls (5.36 and 2.87 vs 6.05 and 3.40 mg glucose eq./100mgDW for the control and infected plants, respectively).

Proline content showed a strong increase on 9 and 23 DAI in the infected plants with respect to the control plants. On 9 DAI, proline levels showed an increase of 48% (Figure 7A) and 17.5% on 23 DAI (Figure 7B). No differences in dry weight content were observed in none of the treatments along the experiments performed (Supplemental Figure 8).

**Figure 7:**
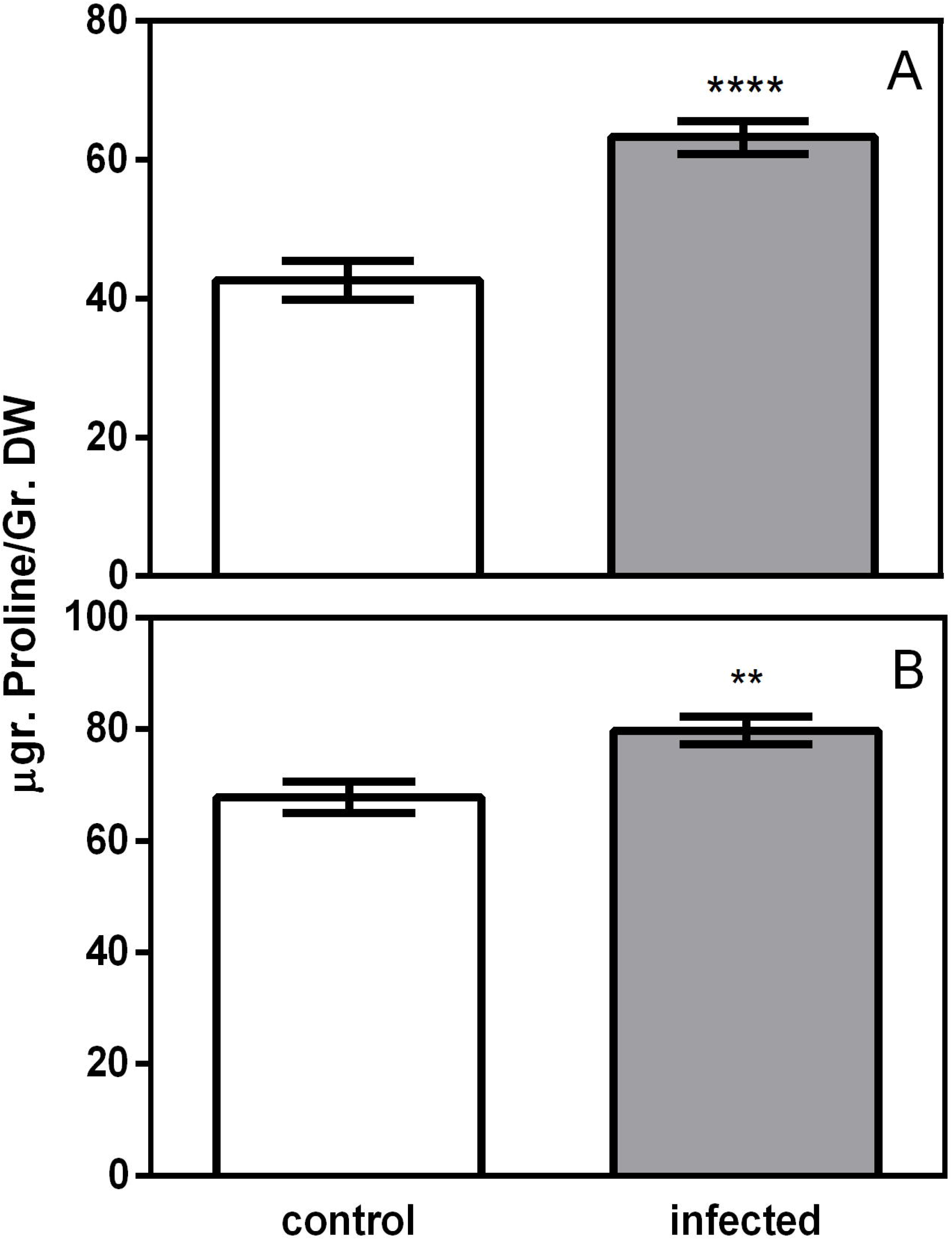
Proline content in leaflets of pecan control plants (white bars) and of infected plants (grey bars) on days 9 (A) and 23 (B) after inoculation. Data are mean with SEM and t-tests were performed for each sampling time comparing control plants and infected plants with a Graph Pad Prism^®^, ns-no significant; * p≤ 0.05; ** p≤ 0.01; *** p≤ 0.0001, n control: 16; n infected: 15.

## 4 DISCUSSION

Pecan plants, like other cultivated plant species, are continuously exposed to potential production constraints which derive from a heterogeneous set of factors or agents that adversely affect pecan nut production, causing significant negative impact in the agricultural industry. Among these potential agents is the dieback which is associated with limb cankers. This complex and pernicious disease which causes dryness and falling branches is caused by *Phomopsis spp.* or by a fungal complex formed by *Phomopsis spp.* and *Botryosphaeria spp.* (Reilly *et al*., 2010). In the present work, we observed typical symptoms of stem canker in plants of pecan cv. Pawnee, artificially inoculated with *Phomopsis spp.* (Figure 1). These symptoms were identical to those previously reported in pecan plants in the southern Pampean region in Argentina (Noelting *et al*., 2016; Mantz personal observation). In this area, *Phomopsis spp.* (anamorphic state of *Diaporthe)* has been identified as the causal agent of cankers in limbs and dieback in branches in many cultivars in a pecan orchard.

According to the mechanism of infection, phytopathogenic fungi are classified into three broad categories, namely necrotrophic, biotrophic, and hemibiotrophic. Necrotrophic fungi kill host cells before colonising them and they can grow and sporulate on dead tissue. In contrast, biotrophic fungi require living host tissue to survive as they feed on nutrients extracted directly from the living cell cytoplasm. Hemibiotrophic pathogens are initially biotrophic but become necrotrophic in a later stage (usually at the spore production stage) (Précigout *et al*., 2020). Taking this classification into account, the majority of *Phomopsis* species are thought to be hemibiotrophic (Udayanga *et al*., 2011). In our study, the symptoms of infection produced by *Phomopsis* were found to be compatible with those resulting from hemibiotrophic infection. The appearance of these symptoms was preceded by an asymptomatic post-inoculation period after which the proper symptoms of the disease emerged in the form of stem cankers.

Plants respond to pathogen attack by inducing the production of myriad proteins and metabolites. When such responses are triggered, Pas levels undergo remarkable changes. Evidence demonstrates that plants use polyamine biosynthetic pathway and oxidative catabolism as a defence mechanism against pathogens. Results from the present study demonstrate that the Pas found in pecan leaves correspond to those that are most abundant in the majority of plants, namely Put, Spd, and Spm (Supplemental Figure 2). In line with this, the free Pas levels measured in pecan leaflets in the present study showed similar values (Figure 2) to those found in other plant species, such as rice seedling leaves (Maiale *et al*., 2004), soybean hypocotyls (Campestre *et al.*, 2011), tobacco and corn seedling (Marina *et al*., 2008, Rodriguez-Kessler *et al*., 2008) and tomato leaves (Vilas *et al*., 2018). Previous studies on Pas content in woody plant species (Bartolini *et al*., 2009; Mirsoleimani and Shahsavar 2018; Liu and Moriguchi 2007; Rey *et al*., 1994) revealed that free total Pas levels ranged from 300 to 430 nmol/grFW in hazelnut leaves, from 500 to 1000 nmol/grFW in malus seedlings, from 437 to 600 nmol/grFW in peach flowers, and from 187 to 477 in nmol/grsFW in Kinnow mandarin leaves. In all these plants, only Put, Spd, and Spm were detected and the values corresponding to the three of them coincided with those reported in fresh weight in the present study. In addition, although the Pas levels in the pecan leaves analysed in the present study were found to be lower than those in the above-mentioned woody plant species, they were in the same range (Figure 3A). Total free Pas levels were observed to decrease over time as leaves aged (Figure 3A) in agreement with observations documented in hazelnut and mandarin leaves (Rey *et al*., 1994, Mirsoleimani and Shahsavar 2018). Further research showed that whereas Pas levels increased in hazelnut trees after severe pruning as an indicator of juvenility and vigour, they decreased in not pruned hazelnut trees, thus indicating ageing and senescence (Rey *et al*., 1994).

Polyamine metabolism in plants infected by pathogens usually varies significantly depending on plant species and pathogen nature. Findings from previous research showed that Pas levels and the activity of polyamine metabolic enzymes are increased in infected tissues during microbial colonisation, which seems to be independent of the pathogenic attack mechanism (Jimenez-Bremont *et al*., 2014). In contrast, findings from the present study indicated a gradual decrease in Put, Spd, and Spm levels in pecan leaves from the infected plants with respect to the controls. In this respect, previous research has shown that in some pathosystems Pas levels increase significantly compared to their controls for a short period of time after which they return either to initial or even lower levels as was observed for example in *Puccinia hordei* - Barley interaction and *Puccinia graminis* - wheat interactions (Greenlands and Lewis 1984, Foster and Walters 1992). In tobacco plants inoculated with *Peronospora tabacina*, *Alternaria tenuis*, *Erysiphe cichoracearum*, *Pseudomonas tabaci*, and tobacco mosaic virus, Pas levels were found to be lower than those in controls (Edreva, 1997). Therefore, taking into account the above-mentioned findings, it appears to be prudent not to discard the possibility that Pas levels increase at shorter times than those evaluated in this work. On the other hand, comparative analyses between tolerant and susceptible cultivars have shown variations in the Pas levels of plant species. In general, cultivars tolerant to a certain pathogen compared to susceptible cultivars have been observed to show a higher increase in Pas (Marini *et al*., 2001, Romero *et al*., 2018). Given this scenario, it is prudent not to discard the possibility that the Pas levels analysed in the present study are due either to a tolerant cultivar or to a low virulent isolate of the pathogen.

Finally, the decrease in Pas levels observed in infected pecan leaves in the present study could be due either to a decrease in the synthesis rate or to an increase in catabolism, including a remobilisation of Pas to other tissues. For example, tobacco plants attacked by the necrotrophic fungus *Sclerotinia sclerotiorum* showed an increase in Put and Spm levels in the apoplast (Marina *et al*., 2008), which was associated to the catabolism of Pas that ends up generating reactive oxygen species and thus collaborating with the death of tissues, which is what is required by this type of pathogen. It was also observed that the addition of Pas to infected tissues increases necrosis (Marina *et al*., 2008).

Transversal cuts in the limb region of *Phomopsis spp.*-infected pecan plants showed dark brown spots in the xylem zone with abundant dead tissue that could accelerate senescence in the leaves (Figure 1B). Senescence has been described as a process associated with a decrease in Pas content (Sobieszczuk-Nowicka *et al*., 2019). In our study, due to the hemibiotrophic nature of *Phomopsis*, a typical pattern of necrotrophic infection was observed after several days post-inoculation, a process that could accelerate mechanisms associated with senescence and consequently modify the levels of Pas.

On the other hand, PSII functionality evaluated by OJIP analysis showed a significant decrease in Sm and ψEO on 23 DAI (Figure 4B, 4D). Sm parameters indicate the number of electron carriers per electron carrier chain, whereas ψEO is the probability that trapped exciton moves an electron into the electron chain beyond QA and represents the energy of the electron transport about the energy trapped (Stirbet and Govindjee, 2011). These data indicate a decrease in the bulk in the electron carriers and a degree of inhibition in the step from QA reduced to QB reduction. At the same time, the QB reducing centre decrease on 23 DAI and the K-band (Woji-Wojc) increase overtime during the experiments (Figure 5) indicated a poor electron transfer from OEC to P680 and showed a similar shape to that produced by DCMU poisoning (Chen *et al*., 2014).

The genus *Diaporthe* is known to produce a set of compounds with antimicrobial and phytotoxic activity (Udayanga *et al*., 2011). Although, *D. helianthi,* in particular, which is the causal agent of stem canker of sunflower, has been reported to produce the phytotoxin Phomozin (Mazars *et al.*, 1990), no reports have been found to date on the effect of this toxin on the PSII. Other authors showed the effect of *D. phaseolorum* methanolic extracts on the PSII activity of *Senecio occidentalis* and *Ipomoea grandifolia* (Moura *et al*., 2020) with PIabs decrease and inhibition in the QB reduction. As the anamorph *Phomopsis* was used in this work, the presence of toxins in diseased plants could be a subject for future research.

Previous research has demonstrated that whereas Pas of three and four amine groups, like Spm and Spd, show photoprotection in isolated thylakoid membranes, Put or methylamine have no effect on this mechanism. This photoprotection is due to the polycationic nature of Spm that stabilises the conformation of PSII protein through electrostatic interaction (Hamdani *et al*., 2011). In the present study, a significant decrease of Spm and Spd was recorded on 23 DAI in the infected plants in agreement with the deterioration of PSII functionality, thus indicating a possible connection between Spm levels and photosynthetic activity which could lead to the hypothesis that *Phomopsis* manipulates Pas metabolism for its benefit.

Sugar concentration is a good indicator of the plant energetic balance during plant-fungus interaction (Nieva *et al*., 2019) and the decay of PSII functionality in infected plants could be associated with lower photosynthetic activity and sugar content. In the present study, both PSII functionality and sugar content were observed to decrease in the infected plants (Supplemental Figure 7B) in agreement with data collected from other plant fungus interactions, such as *Oidium heveae - Hevea brasiliensis*, *Botrytis cinerea - Solanum lycopersicum*, and *Fusarium solani - Lotus tenuis* (Wang *et al*., 2014; Berger *et al*., 2004; Nieva *et al*., 2019).

Furthermore, heat loss occurs via water evaporation which cools the canopy and therefore the subsequent stomatal closure or water supply decrease elicits a leaf temperature increase which could be interpreted as a stress indicator (Costa *et al*., 2013). Pinter *et al*. (1979) found an increase in leaf temperature of 3 to 4 °C in moderately diseased plants of sugar beet and cotton affected by root rot pathogen infection. Other studies have reported similar relationships between pathogen attack to the root system and leaf temperature increase (Tu and Tan 1985; Calderon *et al*., 2013), and further research concluded that vascular damage or obstruction could be a cause of the reduction of water supply to the leaves (Huang *et al*., 2020). In the same direction, the augmented canopy temperature in pecan plants infected with *Phomopsis spp*. and the necrotic xylem tissue in the zone of infection suggest that both events could be related, and further research is necessary to clarify this relationship (Figure 1).

Proline contents increase under different stress types because this amino acid has important roles as membrane structure stabiliser and reactive oxygen scavenger under drought stress (Chun *et al*., 2018). Under mild drought stress, it was observed that whereas proline contents show a significant increase, the relative water content shows no changes, thus suggesting that this amino acid is more sensitive as an indicator of drought stress in pecan leaves (Babuin *et al*., 2016). Taking this into account, the augmented canopy temperature (Figure 6) and the accumulation of proline observed as a consequence of infection in the present study (Figure 7) could be another indicator of drought stress, as documented in pecan plants under restricted water supply (Babuin *et al*., 2016).

Furthermore, whereas the shape of the K-band (Figure 5B) observed in the infected pecan plants coincided with the shape of the K-band in plants with symptoms of drought stress (Oukarroum *et al*., 2007), Abs/RC and DIo/RC increased in the infected plants on 23 DAI in contrast to what occurred in the controls (Supplemental Figure 5) as documented in rubber tree under drought stress (Falqueto *et al*., 2017). All in all, the data collected in the present study suggest that the symptoms resulting from *Phomopsis spp.* infection in pecan plants mimics those derived from drought stress. The physiological effects observed in infected plants in the present study also coincide with symptoms of twig and branch dieback in plants in the field.

As a concluding remark, it can be said that although there are still several unexplored niches, such as −among others− the potential presence of toxins as a result of *Phomopsis spp.* infection in pecan leaves and the effect of this pathogen on stomata regulation and water supply to the leaves, the findings from the present study make an important contribution to understanding the effects of stem canker caused by fungi of the genus of *Phomopsis* on pecan plant physiology.

## Supporting information

SF2: Genealogical Tree constructed with a MEGA 6.06 software using data taken from the GeneBank. The Phomopsis spp. sequence (KU359781.1 MGM1) analyse

SF 3: Polyamine chromatograms obtained in reverse phase HPLC. Standard is a mixture of synthetic polyamines diamopropane, cadaverine, putrescine, hept

SF4: Normalised OJIP curves (Ft-F0)/(Fm-F0) in control plants (black line) and infected plants (grey line) on days 0, 9, 16 and 23 after inoculation.

SF5: Spider plot of a group of parameters obtained via OJIP analysis on days 0 (A), 9 (B), 16 (C) and 23 (D) after inoculation.

SF6: Spider plot of another group of parameters obtained via OJIP analysis on days 0 (A), 9 (B), 16 (C) and 23 (D) after inoculation.

SF7: Glucose (A and C) and starch (B and D) on days 9 (A and B) and 23 (C and D) after inoculation in control plants (white bars) and infected plants

SF8: Dry weight content in leaflets of pecan control plants (white bars) and infected plants (grey bars) on days 9 (A) and 23 (B) after inoculation. D

## Acknowledgements

This work was funded by a grant provided by CONICET (PIP0363-2013). This work is part of the specializing thesis in tree nuts of GMM.

## Data availability statement

The data that support the findings from this study are available from the corresponding authors upon reasonable request.

## Supplemental figure legends

SF 1: Weather data: maximum temperature (black line), minimum temperature (grey line) in ° C, and RH (broken line) in percentage. Arrows indicate the sampling moment. Dots indicate the moment at which canopy temperature was measured.

SF2: Genealogical Tree constructed with a MEGA 6.06 software using data taken from the GeneBank. The *Phomopsis spp.* sequence (KU359781.1 MGM1) analysed in this study was used together with the 28 sequences that showed the greatest similarity in the GeneBank, the sequence MH042300 belonging to *Epicoccum nigrum* strain 180305 was used as a reference.

SF 3: Polyamine chromatograms obtained in reverse phase HPLC. Standard is a mixture of synthetic polyamines diamopropane, cadaverine, putrescine, heptanodiamine, spermidine, and spermine. Control and inoculated chromatograms showed peaks of putrescine, spermidine and spermine, and the internal standard heptanodiamine.

SF4: Normalised OJIP curves (Ft-F_0_)/(Fm-F_0_) in control plants (black line) and infected plants (grey line) on days 0, 9, 16 and 23 after inoculation.

SF7: Glucose (A and C) and starch (B and D) on days 9 (A and B) and 23 (C and D) after inoculation in control plants (white bars) and infected plants (grey bars). Data are mean with SEM and t-tests were performed for each sampling time comparing control plants and infected plants with a Graph Pad Prism^®^. No significant differences were observed.

SF8: Dry weight content in leaflets of pecan control plants (white bars) and infected plants (grey bars) on days 9 (A) and 23 (B) after inoculation. Data are mean with SEM and t-tests were performed for each sampling time comparing control plants and infected plants with a Graph Pad Prism^®^, ns-no significant; * p≤ 0.05; ** p≤ 0.01; *** p≤ 0.0001, n control: 16; n infected: 15.

