## Supplementary figures and images for "Stem canker caused by *Phomopsis spp*. induces changes in polyamine levels and chlorophyll fluorescence parameters in pecan leaves"

### SF2: Genealogical Tree constructed with a MEGA 6.06 software using data taken from the GeneBank. The Phomopsis spp. sequence (KU359781.1 MGM1) analyse

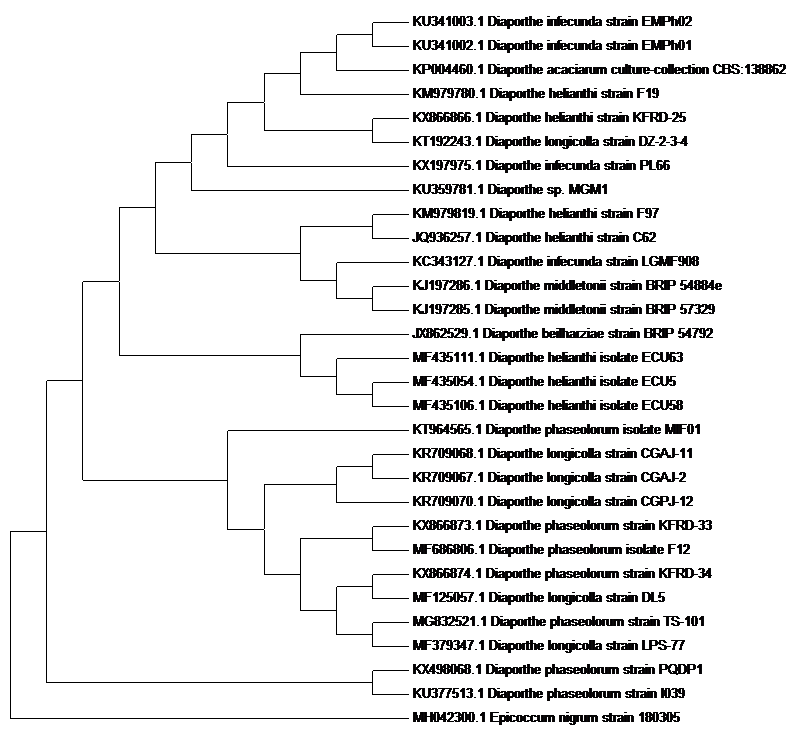

### SF4: Normalised OJIP curves (Ft-F0)/(Fm-F0) in control plants (black line) and infected plants (grey line) on days 0, 9, 16 and 23 after inoculation.

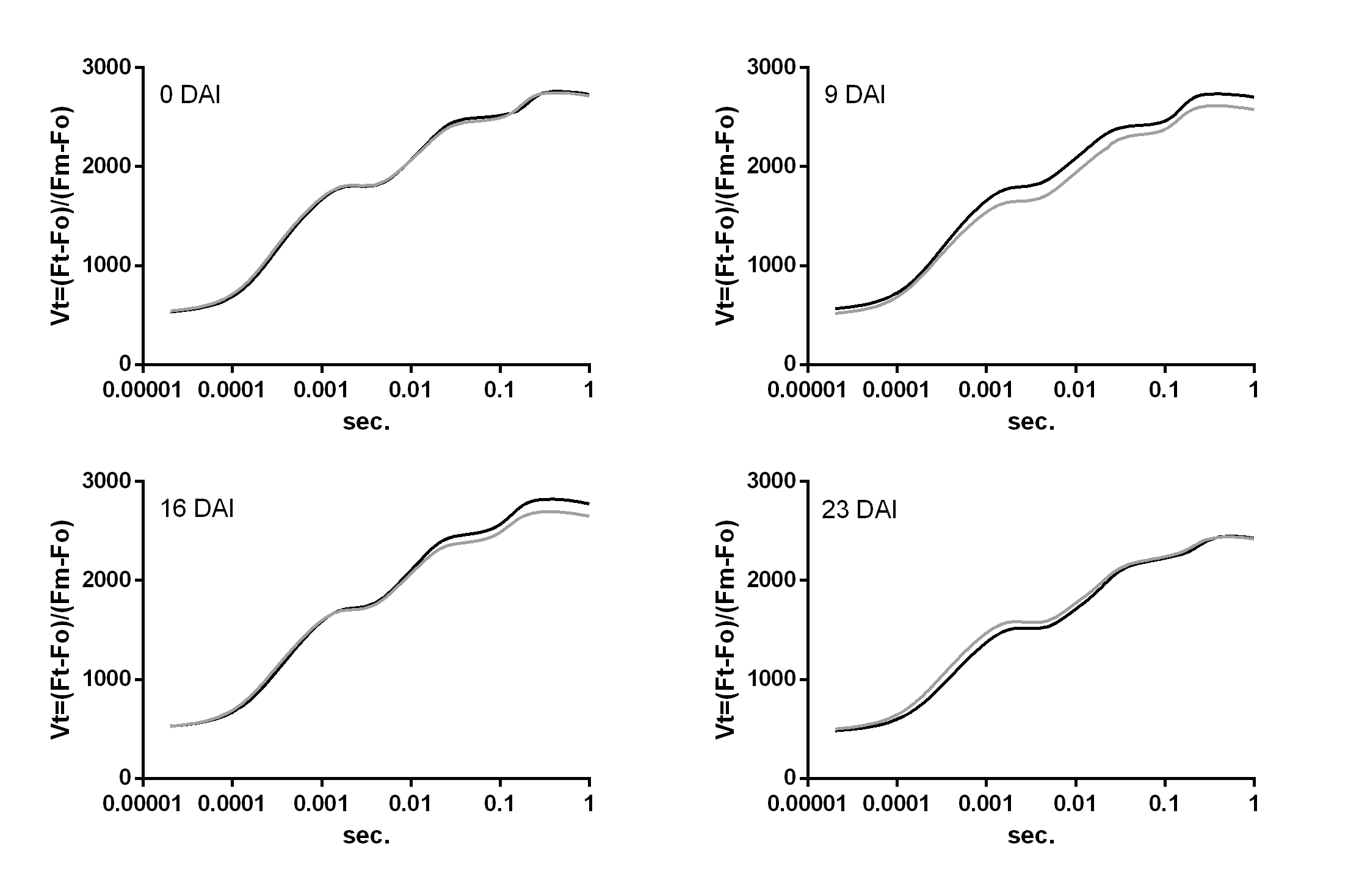

### SF5: Spider plot of a group of parameters obtained via OJIP analysis on days 0 (A), 9 (B), 16 (C) and 23 (D) after inoculation.

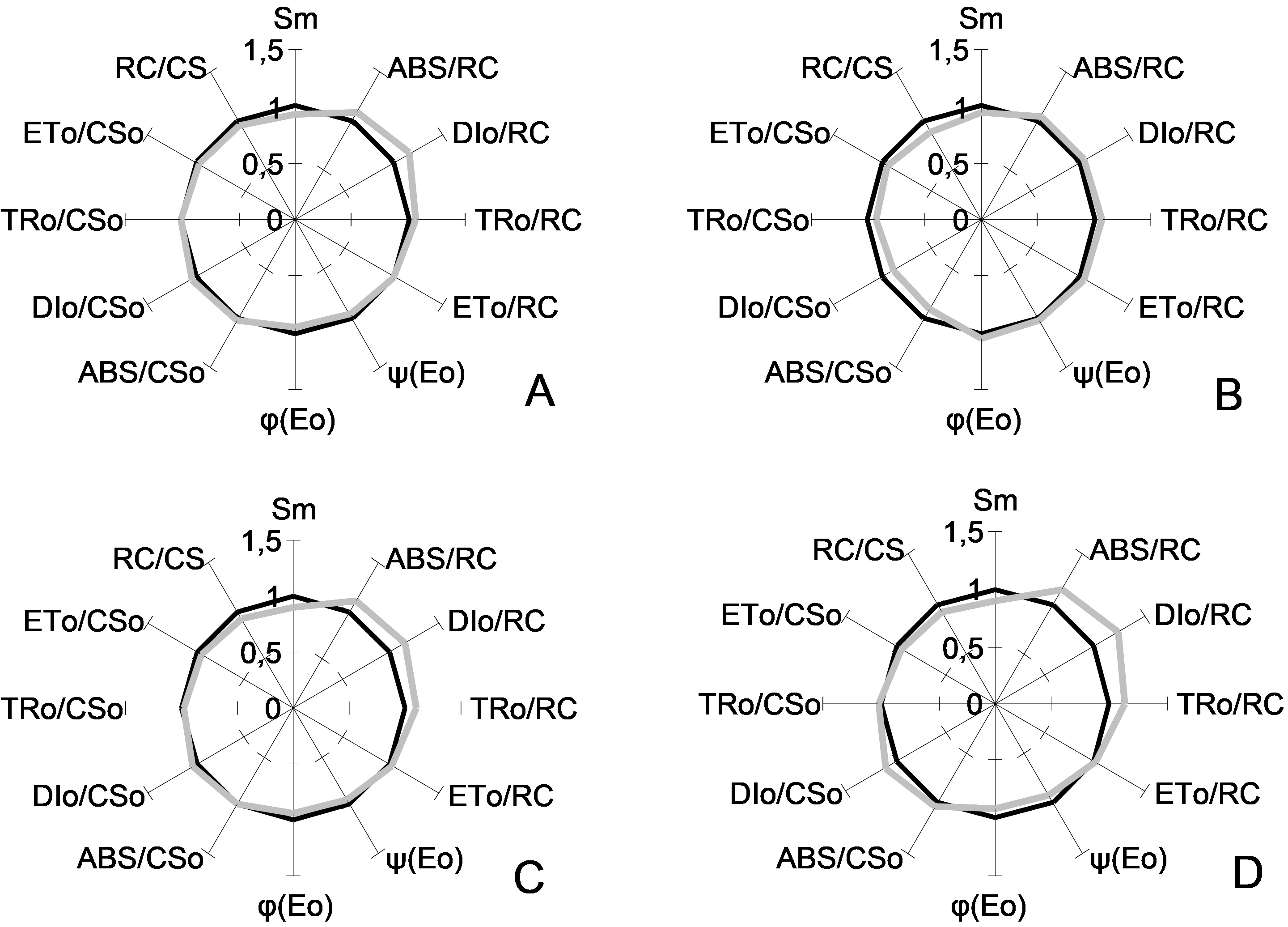

### SF6: Spider plot of another group of parameters obtained via OJIP analysis on days 0 (A), 9 (B), 16 (C) and 23 (D) after inoculation.

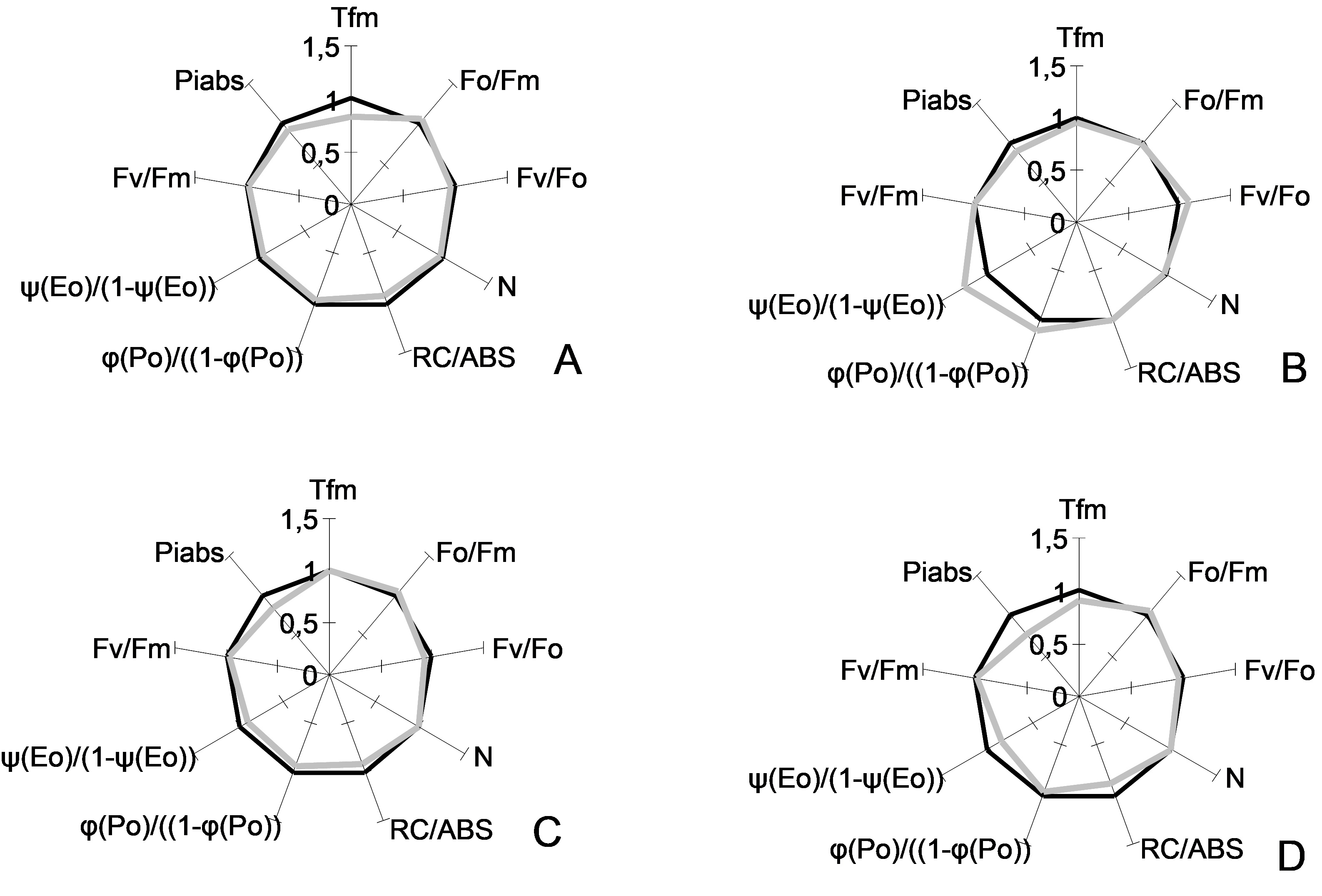

### SF7: Glucose (A and C) and starch (B and D) on days 9 (A and B) and 23 (C and D) after inoculation in control plants (white bars) and infected plants

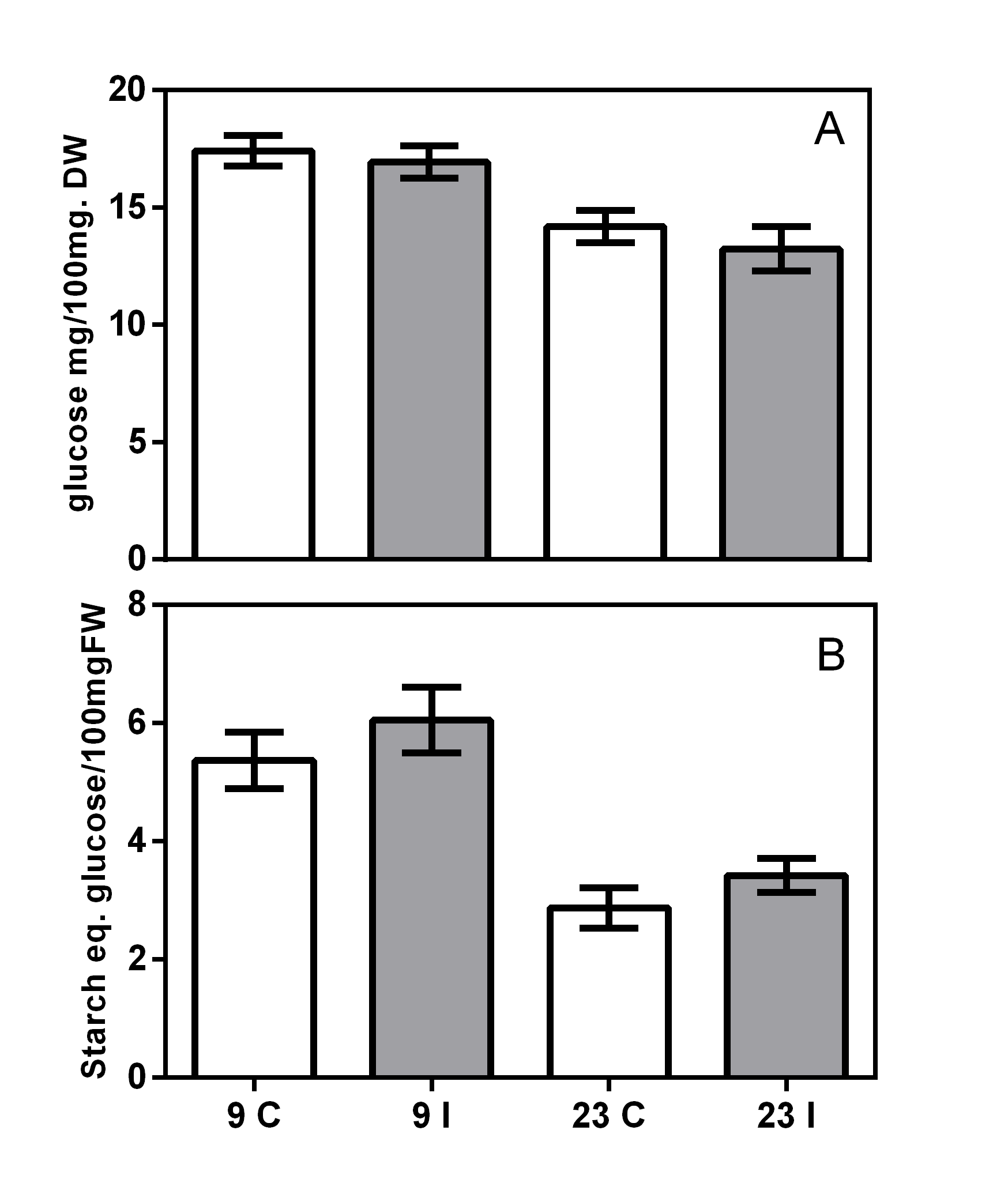

### SF8: Dry weight content in leaflets of pecan control plants (white bars) and infected plants (grey bars) on days 9 (A) and 23 (B) after inoculation. D

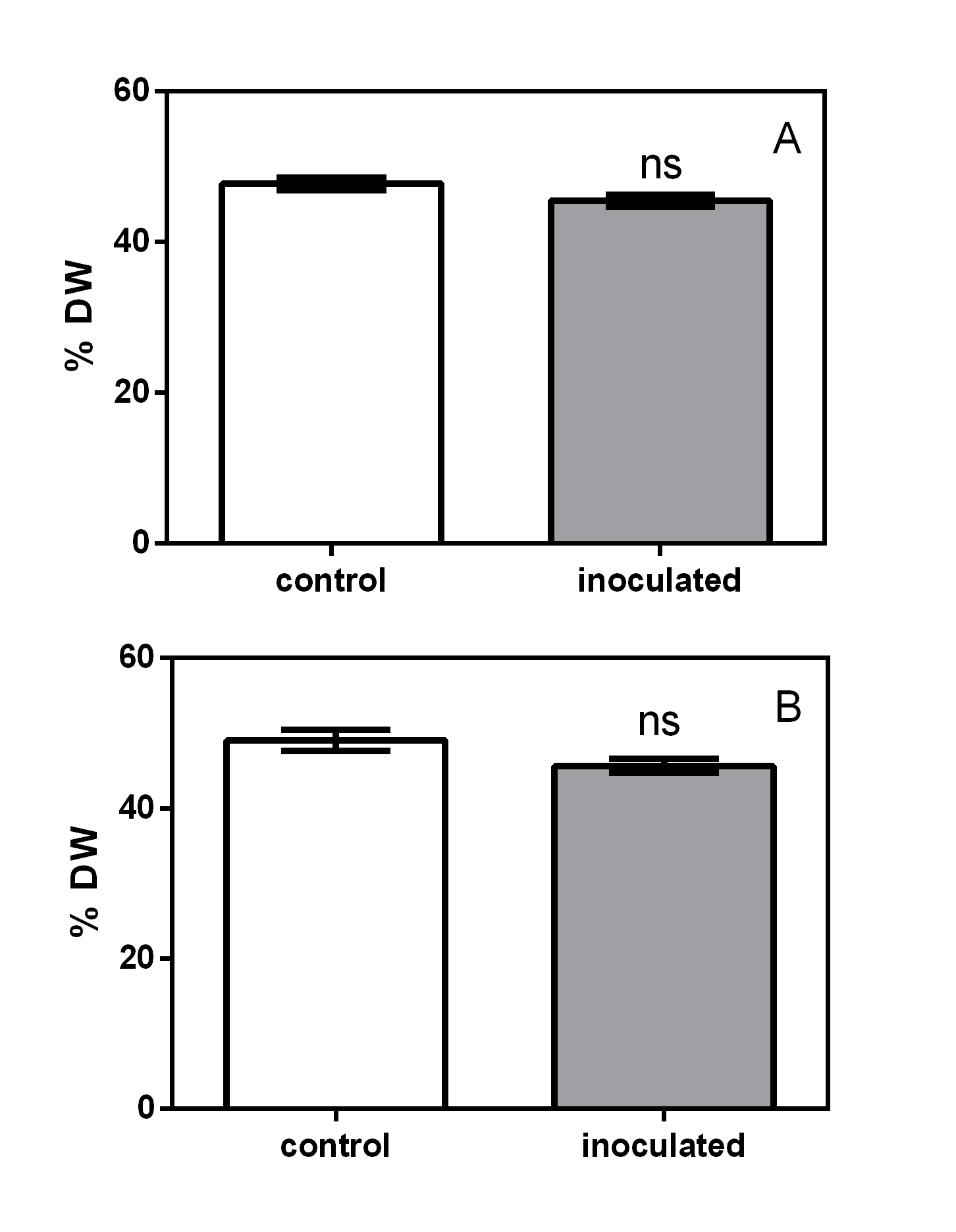

### SF 3: Polyamine chromatograms obtained in reverse phase HPLC. Standard is a mixture of synthetic polyamines diamopropane, cadaverine, putrescine, hept

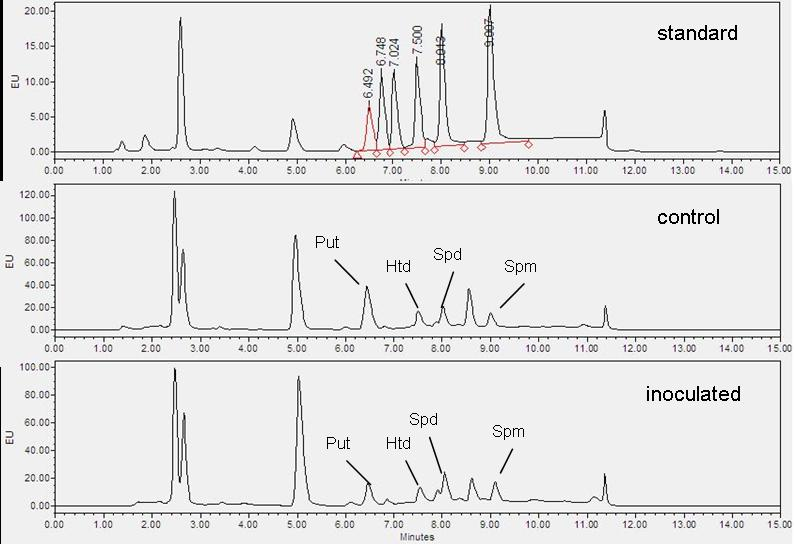

### Supplemental Data 1

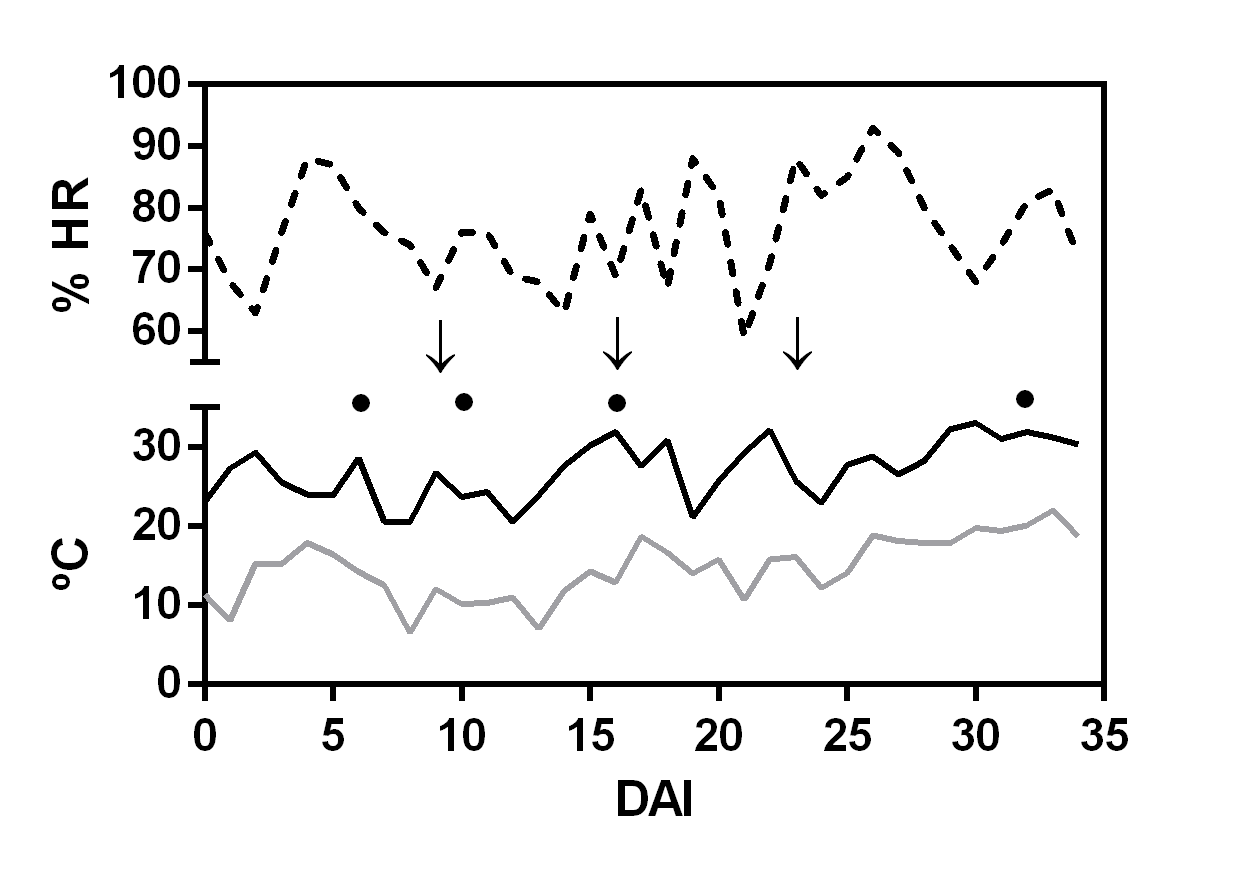
